# Mitochondrial dysfunction triggers secretion of the immunosuppressive factor α-fetoprotein

**DOI:** 10.1101/2021.07.02.450924

**Authors:** Kimberly A. Jett, Zakery N. Baker, Amzad Hossain, Aren Boulet, Paul A. Cobine, Sagnika Ghosh, Philip Ng, Orhan Yilmaz, Kris Barreto, John DeCoteau, Karen Mochoruk, George N. Ioannou, Christopher Savard, Sai Yuan, Christopher Lowden, Byung-Eun Kim, Hai-Ying Mary Cheng, Brendan J. Battersby, Vishal M. Gohil, Scot C. Leary

## Abstract

Signaling circuits crucial to systemic physiology are widespread, yet uncovering their molecular underpinnings remains a barrier to understanding the etiology of many metabolic disorders. Here, we identify a copper-linked signaling circuit activated by disruption of mitochondrial function in the murine liver or heart that results in atrophy of the spleen and thymus and causes a peripheral white blood cell deficiency. We demonstrate that the leukopenia is caused by α-fetoprotein, which requires copper and the cell surface receptor CCR5 to promote white blood cell death. We further show that α-fetoprotein expression is upregulated in several cell types upon inhibition of oxidative phosphorylation, including a muscle cell model of Barth syndrome. Collectively, our data argue that α-fetoprotein secreted by bioenergetically stressed tissue suppresses the immune system, an effect which may explain the recurrent infections that are observed in a subset of mitochondrial diseases or in other disorders with mitochondrial involvement.

## Introduction

To maintain homeostasis, tissue and organ systems must adapt in unison to varying metabolic challenges (Rajan and Perrimon, 2011). These responses involve a cell autonomous component that acts locally at the affected tissue and a complementary, cell non-autonomous component in which secreted factors communicate with other tissues to allow for a coherent systemic response (Taylor et al., 2014, Deng and Haynes, 2016). Therefore, to gain insights into systemic physiology and disease etiology it is imperative that we understand how organs sense their functional state and communicate it to distal tissues (Schinzel and Dillin, 2015, Droujinine and Perrimon, 2016).

A defining feature of human mitochondrial disorders is their tremendous clinical heterogeneity with variable tissue-specificity, onset and severity (Suomalainen and Battersby, 2018). While the importance of oxidative phosphorylation (OXPHOS) to cell fitness is widely recognized, the cell and tissue-specific consequences of the mitochondrial dysregulation that leads to the activation of cellular stress responses that ultimately contribute to disease pathogenesis remain poorly characterized. Therefore, a thorough understanding of these molecular mechanisms and how they contribute to pathogenesis is required. Over the last decade, a compendium of studies in human patients and model organisms points to mitochondrial dysfunction as a potent inducer of the integrated stress response (ISR) (Pakos-Zebrucka et al., 2016). The ISR is an evolutionarily conserved, intracellular signaling cascade that finely tunes cytoplasmic protein synthesis and nuclear gene expression to preserve cellular homeostasis in response to cellular stress (Pakos-Zebrucka et al., 2016). Activating transcription factor 4 (ATF4) integrates the four branches of the ISR, directing the expression and secretion of a number of factors. Two of these branches require the translation initiation factors GCN2 and HRI, and each has been shown to communicate mitochondrial dysfunction (Fessler et al., 2020, Guo et al., 2020, Mick et al., 2020). In the context of OXPHOS defects, the enhanced secretion of GDF15 and FGF21 remodels metabolism and both growth factors are recognized as robust biomarkers for mitochondrial disorders (Lehtonen et al., 2016). However, whether the ISR plays a general role in modulating clinically relevant phenotypes in the molecular pathogenesis of mitochondrial disease remains an outstanding question in the field.

The interplay between mitochondrial dysfunction and the immune system in disease pathogenesis is also poorly understood (DiMauro and Schon, 2003, Parikh et al., 2017, West et al., 2011), yet patients with Barth or Pearson syndrome frequently present with neutropenia or pancytopenia, respectively (Clarke et al., 2013, Manea et al., 2009). A comprehensive study investigating immune function and risk of infection further showed that patients with mitochondrial disease had a higher risk of suffering from recurrent infections (Walker et al., 2014). Here, we establish that an isolated deficiency in cytochrome *c* oxidase (COX) in the liver or heart triggers the secretion of α-fetoprotein (AFP), which then acts systemically to suppress the peripheral immune system by inducing the death of white blood cells (WBCs). AFP therefore appears to play a hitherto underappreciated role in immunomodulatory responses to bioenergetic stress that impact systemic physiology and may contribute significantly to disease progression in patients with mitochondrial disorders who suffer from recurrent infections.

## Results

### Hepatocyte-specific deletion of Sco1 triggers a progressive leukopenia and atrophy of the thymus and spleen

We previously demonstrated that mice lacking the copper chaperone *Sco1* in hepatocytes (*Sco1*^*hep*^) develop normally during the first month of life, but subsequently fail to thrive and have a median life expectancy of 70 days (Hlynialuk et al., 2015). Surprisingly, adult *Sco1*^*hep*^ mice exhibit a severe leukopenia and profound atrophy of the spleen (Hlynialuk et al., 2015); however, it is unclear whether these phenotypes are a direct consequence of ablating *Sco1* expression in hepatocytes or a secondary consequence of liver failure. Therefore, to understand if hepatic deletion of *Sco1* triggers systemic signalling that affects cells and organs of the immune system, we measured complete blood cell (CBC) counts, quantified organ mass and examined the ultrastructure of the thymus in *Sco1*^*hep*^ mice and age-matched littermate controls at post-natal day 18, 27, 37 and 47 (Figure 1). This sampling schedule was selected because at post-natal day 27 (P27) *Sco1*^*hep*^ mice are outwardly indistinguishable from *Control* littermates yet their livers already exhibit a severe, combined COX and copper deficiency (Hlynialuk et al., 2015). We found that the WBC count deficiency in terminal blood drawn from *Sco1*^*hep*^ mice was already manifest at P27 and progressively worsened thereafter (Figure 1A). Notably, we did not detect any changes in red blood cell (RBC) counts (Figure 1A). The onset of the leukopenia preceded any loss in body weight and was ultimately accompanied by the disproportionate atrophy of the spleen and thymus relative to other organs (Figure 1B). These data suggest that the effects on the peripheral immune system in *Sco1*^*hep*^ mice are not secondary to liver failure, and clearly establish that the leukopenia is manifest prior to the atrophy of the thymus and spleen.

**Figure 1.**
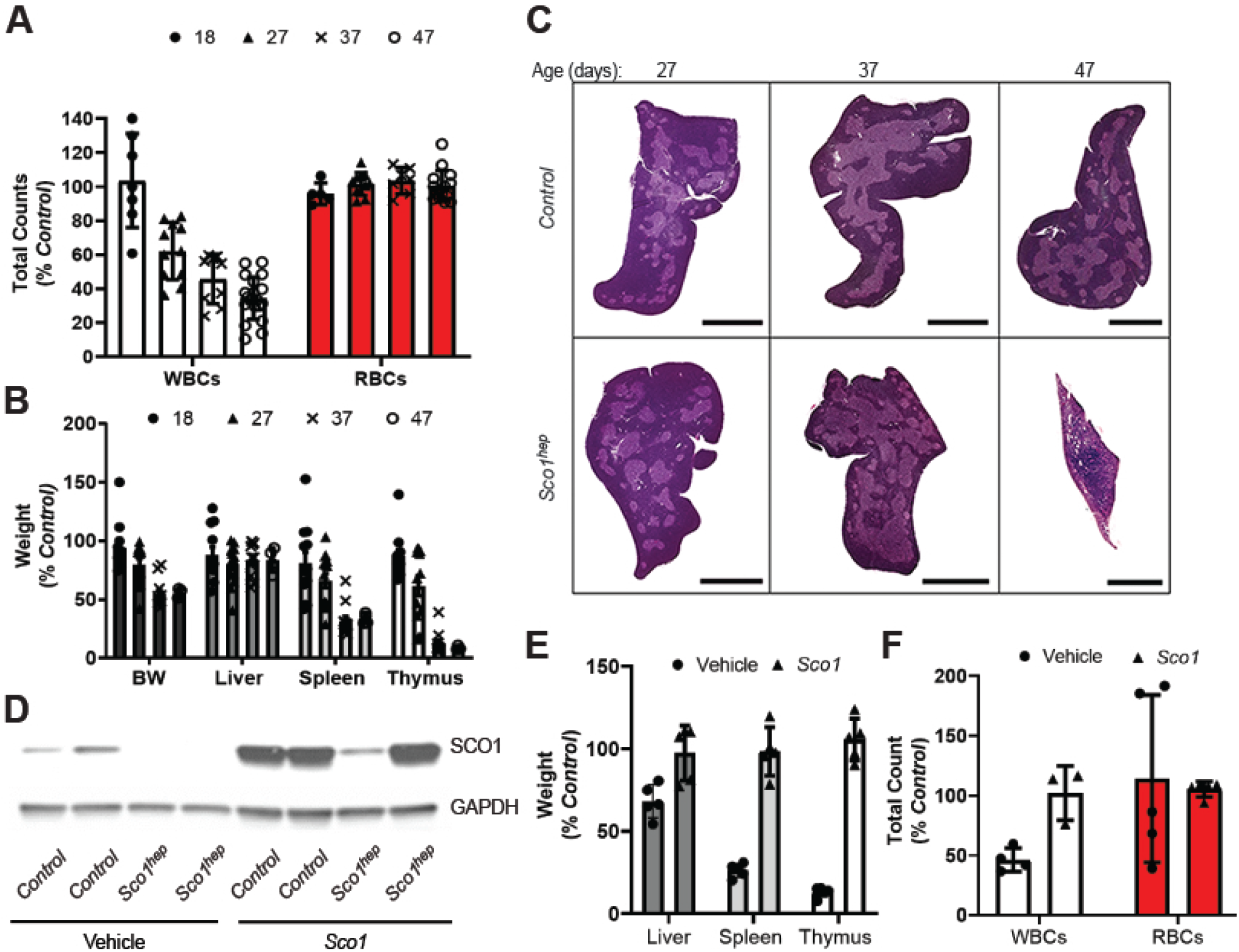
*A)* Progressive leukopenia in *Sco1*^*hep*^ mice (p< 0.01), *B)* disproportionate reduction in the wet weights of the *Sco1*^*hep*^ spleen and thymus at P37 (p<0.01) and P47 (p<0.001) and *C)* selective thinning of the thymic cortex. Scale bars 2mm except for P47 *Sco1*^*hep*^ thymus (800μm). *D)* Adenoviral restoration of SCO1 expression in the liver leads to *E)* rescue of splenic and thymic atrophy and *F)* normalization of WBC counts. Mice were injected at P21 via cardiac puncture with vehicle or helper-dependent adenovirus harbouring a *Sco1* cDNA under the control of a liver-specific promoter and harvested at P47. *Control* refers to wild-type littermates. BW refers to body weight. WBC and RBC denote white and red blood cells, respectively.

To gain further insight into the underlying atrophy of the thymus, we examined thymic ultrastructure using standard hematoxylin and eosin (H&E) staining (Figure 1C). While the thymi of *Control* and *Sco1*^*hep*^ mice are similar at P27, histological analysis of the P37 *Sco1*^*hep*^ thymus revealed a selective thinning of the cortex (Figure 1C) and the presence of tingible body macrophages (Figure S1A). Additional atrophy at P47 was accompanied by a further reduction in the ratio of cortex to medulla, disruption of the cortical-medullary boundary and increased vascularity (Figures 1C, S1A). These data together with our previous findings (Hlynialuk et al., 2015) suggest that loss of *Sco1* function in hepatocytes has a profound effect on the peripheral immune system, and that atrophy of the thymus and spleen in *Sco1*^*hep*^ mice ultimately contributes to the progressive severity of the leukopenia.

### The liver secretes an immunosuppressive factor in response to bioenergetic stress caused by mitochondrial dysfunction

While characterization of the albumin Cre recombinase driver we used for *Sco1* deletion has demonstrated that loxP mediated gene excision is largely restricted to hepatocytes (Postic and Magnuson, 2000), we wanted to ensure that the observed immune phenotypes are indeed directly attributable to the loss of SCO1 function in the liver. We therefore sought to restore *Sco1* expression in the liver of *Sco1*^*hep*^ mice using a helper-dependent adenoviral approach (Brunetti-Pierri and Ng, 2011, Brunetti-Pierri et al., 2013). After establishing that intracardiac administration was superior to intraperitoneal injection for specific delivery of helper-dependent adenovirus to the liver relative to other peripheral organs (Figure S1B), we injected 21-24 day old *Sco1*^*hep*^ mice and *Control* littermates with vehicle or a helper-dependent adenovirus containing a *Sco1* cDNA under the control of a liver-specific PEPCK promoter. SCO1 abundance was significantly increased in the livers of both *Control* and *Sco1*^*hep*^ mice (Figure 1D), which rescued the copper deficiency and the levels of the cellular copper importer CTR1 (Figure S1C, D). Zinc and iron levels were also normalized in the livers of *Sco1*^*hep*^ mice injected with adenovirus (Figure S1C). Critically, adenovirus administration normalized body and organ weights and restored WBC counts in *Sco1*^*hep*^ mice (Figure 1E, F). These data show that the immunosuppressive effects of *Sco1* loss-of-function are specific to its role in hepatocytes and not attributable to aberrant Cre recombinase expression in other cell types.

To determine if the immunosuppressive effect we observe is unique to the *Sco1*^*hep*^ mouse model, we generated two additional hepatocyte-specific knockout lines that lacked the COX assembly factor *Coa5* or *Cox10*. Unlike *Sco1*^*hep*^ mice, and in agreement with existing literature, hepatocyte-specific deletion of *Coa5* or *Cox10* (Diaz et al., 2008) was not lethal (Figure S2A,B). Therefore, *Coa5*^*hep*^ and *Cox10*^*hep*^ mice were sampled at P77 and P64, respectively, as they display the greatest body weight difference at these time points when compared to *Control* littermates (Figure S2A, B). Both *Coa5*^*hep*^ (Figure S2C) and *Cox10*^*hep*^ (Hlynialuk et al., 2015) mice exhibited a severe, combined COX and copper deficiency in the liver, and a significant leukopenia in the absence of changes in RBC counts (Figure 2A, B). Copper levels in the circulation were higher in both *hep* models (Figure S2D), while atrophy of the thymus and spleen was marked in *Coa5*^*hep*^ mice and relatively modest in *Cox10*^*hep*^ animals (Figure 2C).

**Figure 2.**
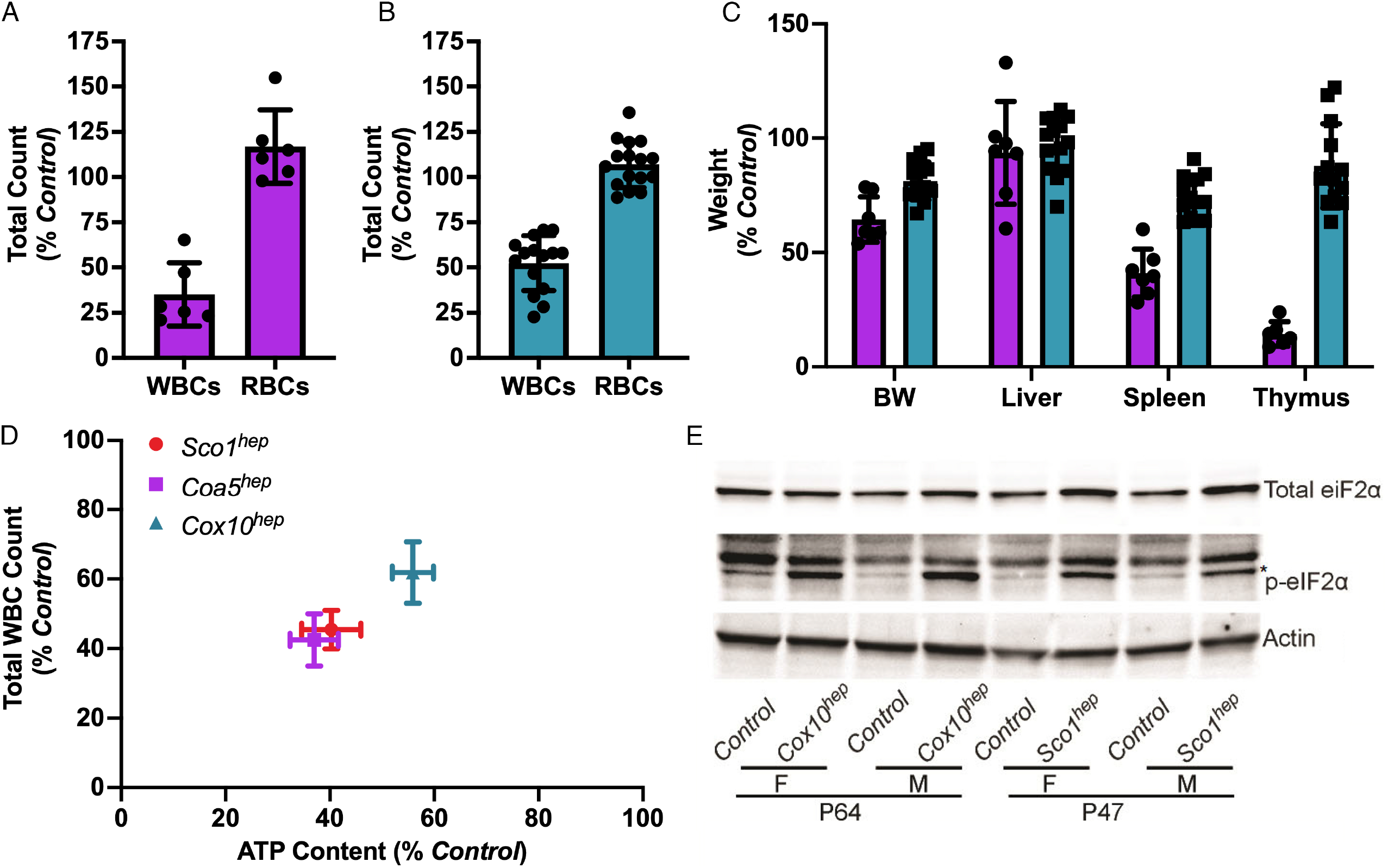
*A) Coa5*^*hep*^ (p<0.001) and *B) Cox10*^*hep*^ (p<0.01) mice also exhibit a significant leukopenia. *C) Coa5*^*hep*^ mice have a disproportionate reduction in the wet weight of the spleen (p<0.01) and thymus (p<0.001), while *Cox10*^*hep*^ mice exhibit significant yet milder atrophy of the spleen (p<0.05). *D)* Total WBC counts are positively correlated with liver ATP content in all three *hep* mouse models (R^2^= 0.99, p=0.001). *E)* P47 *Sco1*^*hep*^ and P64 *Cox10*^*hep*^ livers have higher levels of the phosphorylated form of eIF2α, an ISR marker.

To evaluate whether the varying severity of the leukopenia we observe across our mouse models is attributable to differences in hepatic bioenergetic status, we next quantified total ATP levels in age-matched *Control* and *hep* livers and plotted them against total WBC counts. These analyses revealed a significant, positive correlation between hepatic ATP content and WBC counts (Figure 2D). Consistent with the idea that perturbed energy homeostasis is associated with mitochondrial dysfunction, *Sco1*^*hep*^ and *Cox10*^*hep*^ livers exhibit a significant increase in the abundance of the phosphorylated form of eIF2α, a key player in the ISR (Harding et al., 2003, Pathak et al., 1988) (Figure 2E).

Taken together, our findings from three unique mouse models argue that mitochondrial dysfunction in the liver triggers the secretion of a factor that suppresses the peripheral immune system. Our data further suggest that the severity of the leukopenia is a direct reflection of the extent to which the *hep* liver is bioenergetically stressed as a result of impaired mitochondrial function.

### Hepatic mitochondrial dysfunction leads to the secretion of the immunosuppressive factor AFP

To establish that a secreted factor is responsible for the immune phenotypes we observe as a result of mitochondrial dysfunction in the liver, we serially administered *Control* or *Sco1*^*hep*^ plasma to 21-24 day old *Control* mice over 28 days via tail vein injection. Mice injected with *Sco1*^*hep*^ plasma exhibited a significant leukopenia relative to those injected with *Control* plasma (Figure 3A) independent of alterations in thymic mass (Figure S3A), emphasizing that the leukopenia in our *hep* models does not require atrophy of the spleen or thymus. Taken together, these findings are consistent with a model whereby the *Sco1*^*hep*^ liver is secreting a factor into the blood that is suppressing the peripheral immune system.

**Figure 3.**
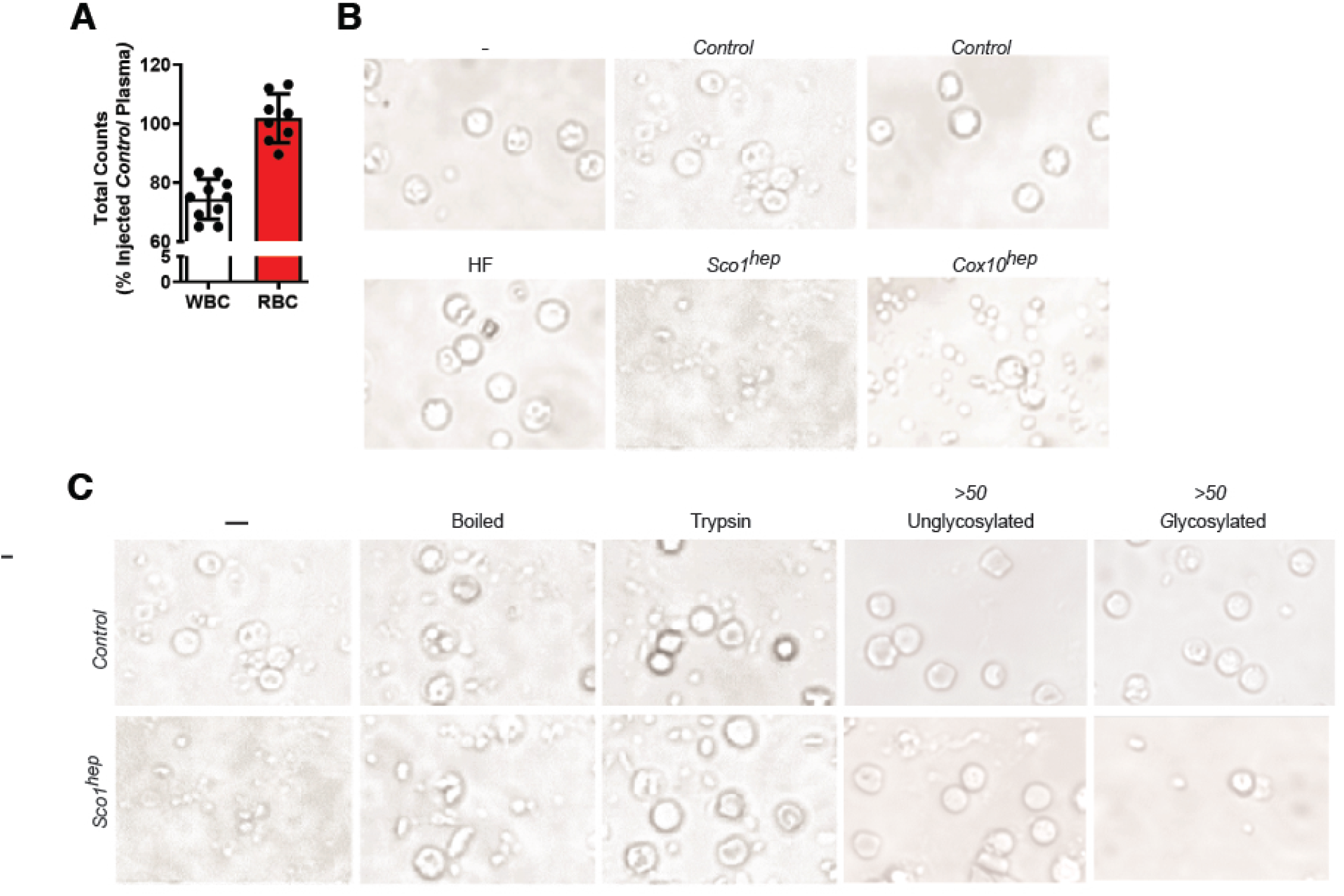
*A)* Total WBC counts are reduced in *Control* mice injected with *Sco1*^*hep*^ plasma relative to those injected with *Control* plasma (p=0.02). *B)* The viability of PBMCs isolated from *Control* mice is reduced when they are co-cultured with *Sco1*^*hep*^ or *Cox10*^*hep*^ plasma. PBMC viability is unaffected by exposure to plasma from mice fed a high fat (H*F*) diet. *C)* PBMC viability is rescued if *Sco1*^*hep*^ plasma is boiled or treated with trypsin. Fractionation based on size and the presence of a glycan revealed that the factor(s) that reduces PBMC viability is contained in a >50kDa glycoprotein fraction. For *B)* and *C)* the – denotes PBMCs cultured in FBS alone.

A number of secreted molecules including proteins, lipids, nucleic acids and metabolites could account for the observed immune phenotypes (Krysko et al., 2011, Kim et al., 2017). To distinguish between these possibilities, we performed an *in vitro* primary blood mononuclear cell (PBMC) viability assay with *Control* and *hep* plasma sources. PBMCs grown in RPMI containing *Control* plasma are indistinguishable from those grown in media supplemented with fetal bovine serum (Figures 3B, S3D; -vs *Control*). In contrast, the viability of PBMCs co-cultured with *Sco1*^*hep*^ or *Cox10*^*hep*^ plasma was markedly diminished (Figures 3B, S3D). Because *Sco1*^*hep*^ livers have profound alterations in lipid metabolism reminiscent of non-alcoholic fatty liver disease (NAFLD) (Hlynialuk et al., 2015), we next addressed the possibility that the observed effects in PBMC cultures were attributable to one or more lipid species. We found that PBMC viability was unaffected by co-culture with plasma isolated from mice fed a high fat (HF) diet (Figures 3B, S3D) (Savard et al., 2013) that had fatty livers without any discernible changes in their hepatic metal ion content or the abundance of core subunits of each OXPHOS enzyme complex (Figure S3B, C). In contrast, boiling or trypsin treatment of *Sco1*^*hep*^ plasma abolished its negative effect on PBMC viability (Figure 3C), suggesting that the secreted factor is a protein. We therefore fractionated plasma proteins based on size and the presence of a glycan, and found that the bioactivity was retained in a >50 kDa glycoprotein-containing fraction (Figure 3C).

To identify the secreted factor, the >50kDa glycoprotein-containing *Control, hep* and *HF* plasma fractions were analyzed by quantitative mass spectrometry (MS) (Figure 4A). We reasoned that the immunosuppressive factor would be absent in *HF* plasma, enriched in *hep* plasma and more abundant in *Sco1*^*hep*^ than *Cox10*^*hep*^ plasma, given the relative severity of the leukopenia in each model. Of the two up-regulated hits, only AFP met those criteria (Figures 4A, S4A). Consistent with our MS results, AFP abundance was elevated in P47 *Sco1*^*hep*^, P77 *Coa5*^*hep*^ and P64 *Cox10*^*hep*^ plasma (Figure 4B), and its circulating levels were significantly higher in *Sco1*^*hep*^ than *Control* plasma at P27 and continued to increase as the severity of the leukopenia worsened over time (Figure 4C). In fact, the roughly 2000-fold difference in AFP abundance between *Control* and *Sco1*^*hep*^ plasma in P47 mice was mirrored at the transcript level in the *Sco1*^*hep*^ liver (Figure S4B), emphasizing that the liver is the principal organ responsible for secreting AFP into the circulation.

**Figure 4.**
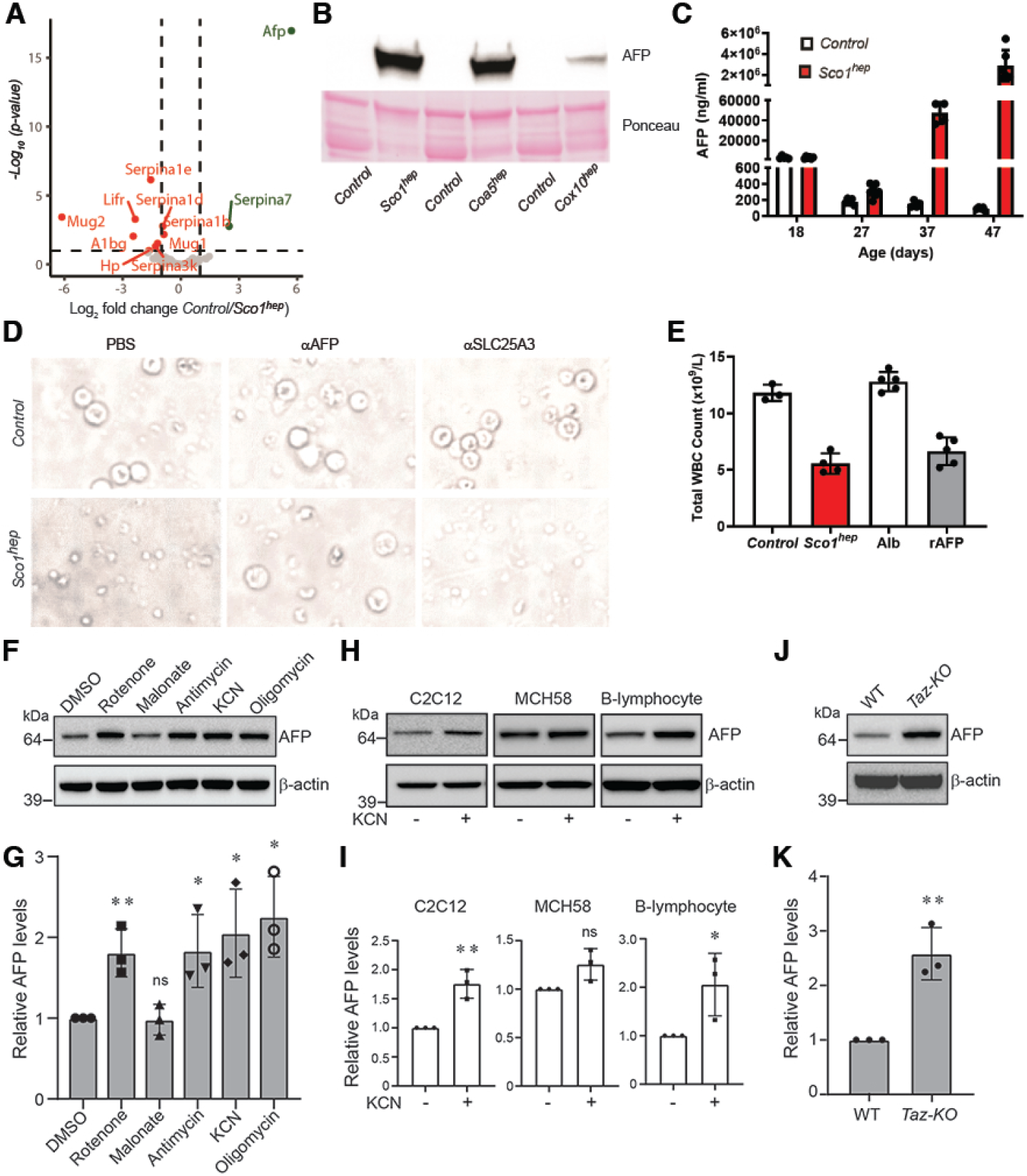
*A)* AFP is significantly enriched in *hep* relative to *Control* plasma. Volcano plot with dotted lines indicating a 2-fold change and adjusted p-value significance threshold of 0.1. A green or red symbol indicates a protein whose abundance is significantly up- or down-regulated, respectively. Grey symbols denote proteins whose abundance is not significantly different (ns) between the two groups. *B)* AFP abundance is markedly upregulated in *Sco1*^*hep*^, *Coa5*^*hep*^ and *Cox10*^*hep*^ plasma. Plasma was pooled (minimum of two males and two females per pool) and depleted of immunoglobulins and albumin prior to Western blotting. The Ponceau stained membrane indicates relative loading across lanes. *C)* AFP progressively accumulates in *Sco1*^*hep*^ plasma (P27, p< 0.05, P37, p< 0.0001, P47, p< 0.0001). *D)* PBMC viability is rescued by immunodepleting AFP from *Sco1*^*hep*^ plasma. Culture media supplemented with phosphate buffered saline (PBS) and plasma incubated with α SLC25A3, an antibody isotype control, served as internal controls. *E) Control* mice develop a leukopenia following serial injection with *Sco1*^*hep*^ plasma (p<0.01) or 1μg of recombinant AFP (rAFP) (p<0.01). *Control* mice injected with *Control* plasma or albumin served as internal controls. *F&G)* Inhibition of the mitochondrial respiratory chain elevates AFP abundance in C2C12 myoblasts. *H&I)* AFP levels increase in C2C12 myoblasts and human B-lymphoblasts upon inhibition of COX *J&K)* Deletion of *Tafazzin* (*Taz-KO*) in C2C12 myoblasts results in higher steady-state levels of AFP. WT refers to wild-type. For panels *F-K)* β-actin was used as a loading conrol and data are shown as mean ± SD, n=3. \**p* < 0.05, \*\**p* < 0.005. ns, not significant. \*\**p* < 0.005.

To further validate AFP as the active component of our signaling axis, we investigated the effect of manipulating its abundance on WBC viability both *in vitro* and *in vivo*. The viability of PBMCs co-cultured in *Sco1*^*hep*^ plasma was rescued upon immunodepletion of AFP but not by pre-treatment of plasma with anti-SLC25A3, an isotype antibody control (Figure 4D). Supplementation of standard media with recombinant AFP (rAFP) alone also induced PBMC death (Figure S4C). Corroborating our *in vitro* findings, serial injection of 21-24 day old *Control* mice with rAFP, but not with the closely related family member albumin, resulted in a leukopenia of comparable severity to that seen in *Control* mice injected with *Sco1*^*hep*^ plasma (Figure 4E). These data collectively argue that AFP secreted by the liver is directly responsible for the unexpected immunosuppression we observe in our *hep* mouse models of mitochondrial disease.

### AFP can be produced by non-hepatic models of mitochondrial dysfunction and requires copper for its immunosuppressive activity

AFP is highly expressed during embryonic development and is also secreted by the yolk sac, stomach and cells of the intestine (Jones et al., 2001). While its expression is repressed post-birth by epigenetic mechanisms (Belayew and Tilghman, 1982, Vacher and Tilghman, 1990), it is known that the adult liver re-activates the *AFP* locus and secretes the protein in response to fibrosis, hepatitis and several hepatic cancers (Abelev, 1971, Castaneda and Pearce, 2008, Nakano et al., 2017). We therefore investigated whether mitochondrial dysfunction is a general trigger for AFP overexpression in other cell types. To test this idea, we treated the murine myoblast C2C12 cell line with pharmacological agents to inhibit specific OXPHOS complexes. Indeed, specific inhibition of COX (Complex IV) and Complexes I, III or V resulted in a modest but significant increase in steady-state AFP levels (Figure 4F, G). To determine whether this is a cell-type dependent phenotype, we treated three additional cell types with cyanide and found that skeletal myoblasts and B-lymphocytes, but not fibroblasts, up-regulated AFP production in response to COX inhibition (Figure 4H, I). These data suggest that in addition to hepatocytes, several other cell types are able to increase AFP production in response to mitochondrial dysfunction.

To expand upon our pharmacological analyses, we investigated whether AFP expression is upregulated in a non-hepatic mitochondrial disease model. We chose to use murine C2C12 skeletal muscle cells that lack *TAFAZZIN* (TAZ) (Lou et al., 2018), the cardiolipin remodelling enzyme whose loss-of-function results in Barth syndrome (Ghosh et al., 2019). Our selection of a Barth syndrome model was guided by the observation that approximately 90% of Barth syndrome patients exhibit persistent or intermittent neutropenia in addition to skeletal muscle and cardiac myopathy (Clarke et al., 2013). We found that AFP abundance was ∼2.5 fold higher in TAZ-KO C2C12 myoblasts compared to isogenic controls (Figure 4J, K). Consistent with this observation, COX-deficient murine hearts lacking the high affinity copper importer CTR1 (Kim et al., 2010) have significantly higher levels of *Afp* transcript (Figure S5A) and protein (Figure S5B) and elevated levels of the ISR marker phospho-eIF2α (Figure S5B) when compared to *Control* littermates. Taken together, these findings indicate that mitochondrial dysfunction triggers re-activation of AFP expression in diverse cell types and tissues.

Several proteoforms of AFP are known to be present in the general circulation and at least some of these have the ability to bind a variety of ligands, including copper (Aoyagi et al., 1978). Therefore, an intriguing possibility is that the leukopenia in *hep* mice is caused by a specific conformer(s) constituting a fraction of the total plasma AFP pool. To address this possibility, we took further advantage of the *Ctr1*^*hrt*^ mouse model because its COX-deficient heart is known to communicate with the liver leading to a significant hepatic copper deficiency (Kim et al., 2010). Like several of our *hep* models, P10 *Ctr1*^*hrt*^ mice exhibited disproportionate atrophy of the spleen and thymus (Figure 5A) and a significant leukopenia (Figure 5B). Due to their young age, circulating AFP levels were comparable in *Control* and *Ctr1*^*hrt*^ animals (Figure 5C); however, only *Ctr1*^*hrt*^ plasma was capable of killing PBMCs (Figure 5D). This observation is consistent with the idea that AFP abundance alone does not explain its immunosuppressive properties. In fact, *Ctr1*^*hrt*^ and *Sco1*^*hep*^ plasma pre-treated with the Cu(I)-specific chelator bathocuproine disulfonic acid (BCS) prevented the induction of PBMC death (Figure 5D, E) whereas the addition of copper salts to the culture media enabled the AFP-rich, age-matched (P10) *Control* plasma to stimulate PBMC death (Figure 5D). Collectively, these data argue that AFP requires copper to induce cell death in WBCs and cause a leukopenia.

**Figure 5.**
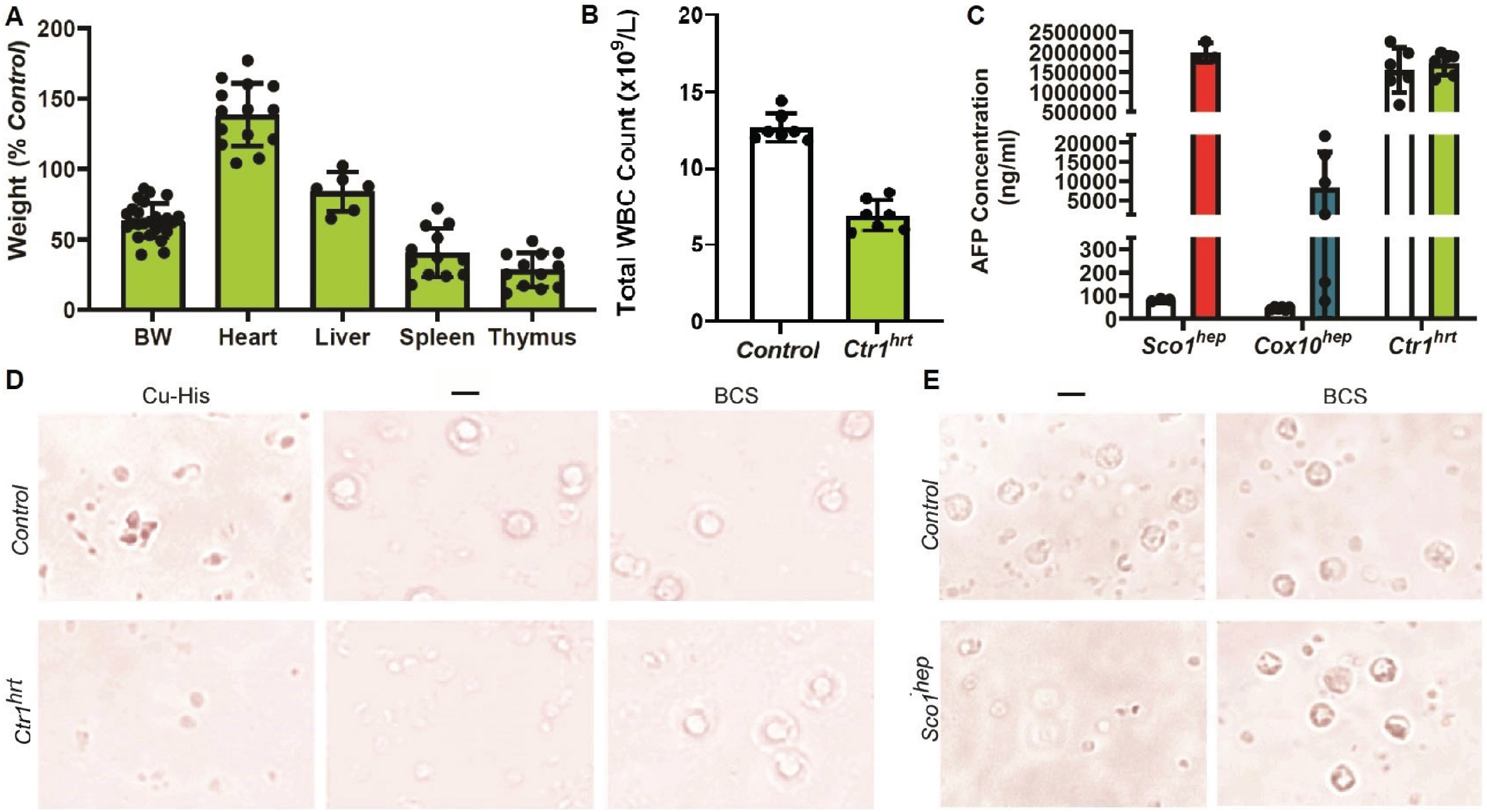
*A&B) Ctr1*^*hrt*^ mice exhibit disproportionate atrophy of the spleen (p<0.01) and thymus (p< 0.01) and have a *B)* leukopenia (p< 0.01). *C)* Plasma AFP levels are comparable in P10 *Control* and *Ctr1*^*hrt*^ plasma mice. Relative levels of AFP in *Sco1*^*hep*^, *Cox10*^*hep*^ age-matched littermate *Control* plasma were quantified at the same time and are shown here for comparative purposes. *D)* PBMC viability is adversely affected by *Ctr1*^*hrt*^ plasma, an effect that can be rescued by adding the copper chelator bathocuproine sulphonate (BCS) to the media. Viability of PBMCs is also reduced if copper-histidine (Cu-His) is added to media containing P10 *Control* plasma. *E)* BCS also negates the negative effect of *Sco1*^*hep*^ plasma on PBMC viability.

### AFP promotes the death of mouse and human PBMCs via the CCR5 receptor

To establish that AFP causes a leukopenia by stimulating WBC death, we isolated PBMCs from P47 *Control* and *Sco1*^*hep*^ mice and cells positive for CD44, a marker of activation, and annexin V, an indicator of cell death. Both markers were significantly elevated in *Sco1*^*hep*^ PBMCs when compared to *Control* PBMCs (Figures 6A, S6), consistent with the idea that AFP indeed stimulates WBC death. To confirm our *in vivo* observations and further delineate the temporal relationship between activation and cell death in the presence of bioactive AFP, we conducted a time course experiment using naïve, wild-type PBMCs co-cultured in *Control* or *Sco1*^*hep*^ plasma. At 12 hours, CD44 staining was significantly higher in PBMCs grown in media containing *Sco1*^*hep*^ plasma while annexin V staining was similar in *Control* and *Sco1*^*hep*^ co-cultures (Figure 6B, C). Annexin V positive cell numbers then increased significantly in the *Sco1*^*hep*^ co-culture over the next 36 hours (Figure 6B), indicating that activated PBMCs were indeed dying.

**Figure 6.**
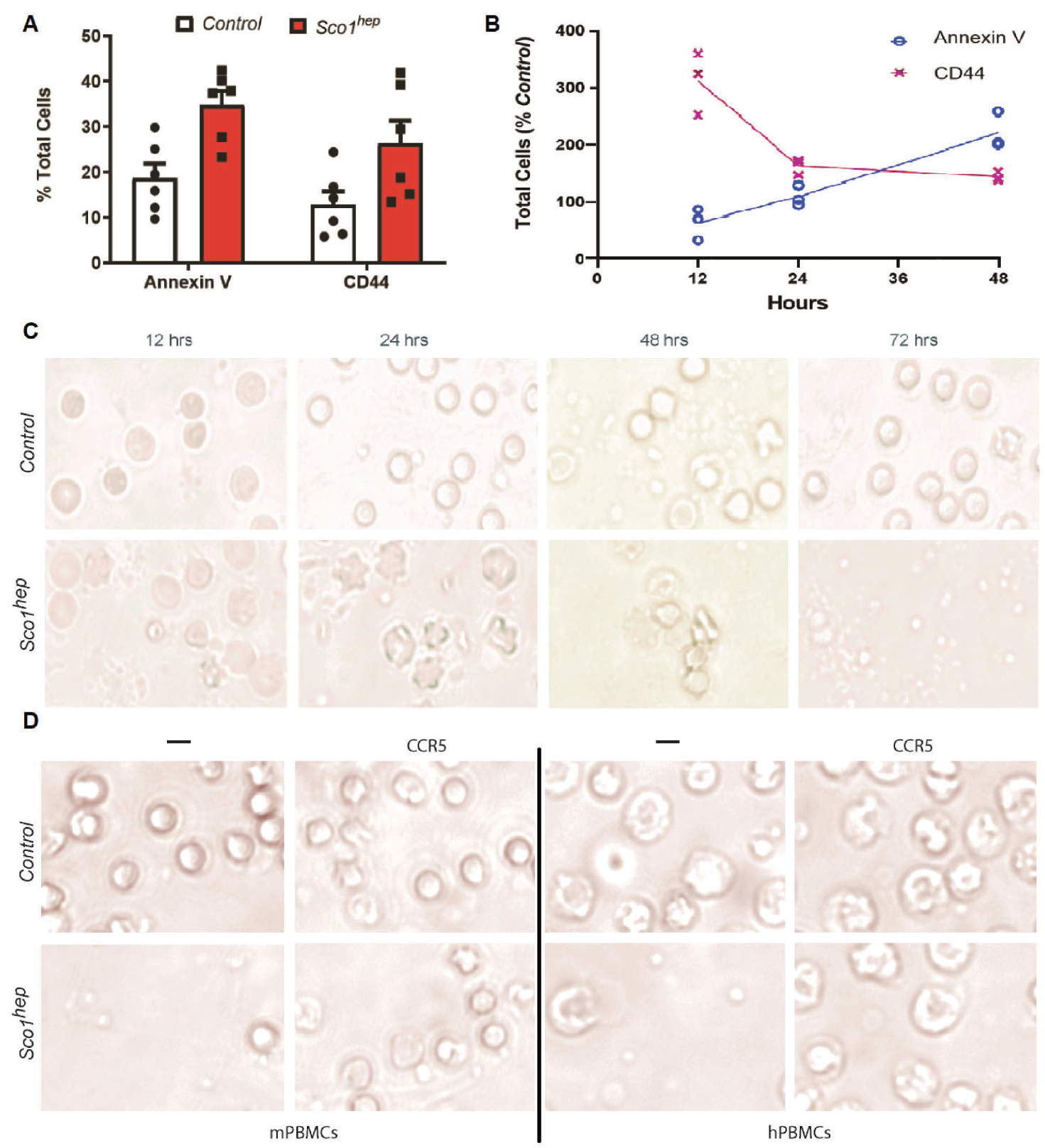
*A)* PBMCs isolated from peripheral blood of *Sco1*^*hep*^ mice have an increased number of cells that stain positive for the activation marker CD44 and the apoptotic marker Annexin V when compared to PBMCs isolated from *Controls. B)* PBMCs isolated from *Control* mice were co-cultured in media containing *Sco1*^*hep*^ plasma are activated earlier and demonstrate a progressive increase in cell surface expression of the apoptotic marker annexin V relative to PBMCs cultured in *Control* plasma. *C)* Cells analyzed in *B)* show blebbing (a sign of apoptosis) as early as 12-24 hrs in culture and loss of cellularity after 72 hr in culture in response to co-culture with *Sco1*^*hep*^ but not *Control* plasma. *D)* Human PBMC viability is also reduced in media containing *Sco1*^*hep*^ plasma, and can be rescued with the CCR5 antagonist Maraviroc.mPBMCs and hPBMCs denote PBMCs of mouse or human origina, respectively.

It has been proposed that AFP exerts its immunosuppressive effects by binding to various cation channels and classes of receptors that include mucin, scavenger, lysophospholipid and chemokine receptors (Zhu et al., 2018, Newby et al., 2005, Mizejewski, 2017, Mizejewski, 2015, Mizejewski, 2014, Mizejewski, 2011). To date, however, AFP has only been shown to bind directly to the chemokine receptor CCR5 (Atemezem et al., 2002). We therefore co-cultured PBMCs in *Control* or *Sco1*^*hep*^ plasma alone or in media that also contained the CCR5 antagonist Maraviroc (Dorr et al., 2005) and found that receptor blocking rescued cell viability (Figure 6D). To determine if AFP from our *hep* mouse models of mitochondrial disease also acts in an immunosuppressive manner on human cells via the same mechanism, we repeated the experiment and found that pharmacologically blocking the CCR5 receptor ablated the ability of *Sco1*^*hep*^ plasma to kill human PBMCs (Figure 6D). Taken together, our data argue that AFP requires both copper and CCR5 to activate and kill WBCs to induce a leukopenia. Our findings further establish that this immunosuppressive mechanism is also capable of triggering the death of human PBMCs and suggest that tissue release of a specific AFP conformer in response to mitochondrial dysfunction provides a mechanism to suppress the peripheral immune system.

## Discussion

Here, we uncover a molecular mechanism by which a primary mitochondrial dysfunction in the liver compromises the peripheral immune system, revealing the basis of a surprising inter-tissue signaling circuit. We demonstrate that a defect in OXPHOS in the post-natal mouse liver induces robust secretion of AFP which, in concert with copper, activates cell death in leukocytes by interacting with the cell surface receptor CCR5. In turn, this leads to a progressive leukopenia and atrophy of the thymus and spleen. Patients with inherited mitochondrial disorders are well-known to be susceptible to acute febrile infections (Muraresku et al., 2018) and in some cases manifest with congenital neutropenia (Spoor et al., 2019); however, a molecular basis for these phenotypes has been lacking until now. Together, our findings identify an immunosuppressive mechanism that could account for why some groups of patients with mitochondrial dysfunction are vulnerable to infections. It may also explain why patients with elevated circulating AFP levels owing to ataxia telangiectasia or other cerebellar ataxias with suspected mitochondrial involvement are also prone to recurrent infection (Waldmann and McIntire, 1972, Renaud et al., 2020). However, considering the tremendous clinical heterogeneity of mitochondrial diseases it is likely that additional mechanisms contribute to immunosuppression in a context-dependent manner that have yet to be identified.

AFP is a major serum protein expressed by the developing embryo that is under tight temporal and spatial regulation (Belayew and Tilghman, 1982, Krumlauf et al., 1986). Post-birth, multiple genetic factors coordinate a conserved developmental program to repress transcription via elements upstream of the gene promoter (Perincheri et al., 2005, Vacher and Tilghman, 1990, Xie et al., 2008). However, liver dysfunction in adults can reactivate AFP expression in hepatocytes, most notably with hepatic cellular carcinoma (HCC) and fibrosis (Abelev, 1971, Nakano et al., 2017). Though the precise functional consequence of AFP secretion in these disease settings is poorly understood, it has been suggested to increase hepatocyte cell proliferation in HCC. In the context of mitochondrial dysfunction, hyperplasia of hepatocytes is not observed (Hlynialuk et al., 2015) nor are high circulating levels of AFP alone toxic to leukocytes. Instead, the leukocyte toxicity requires AFP and copper, a known ligand of this serum protein (Aoyagi et al., 1978).

Copper is an essential metal that acts as a co-factor for select cellular enzymes catalyzing redox reactions. This trace element is obtained from the diet and then systemically distributed to meet cellular demand. In mammals, the liver acts as the principal storage tissue to coordinate systemic copper levels, a function that is of utmost importance in early post-natal life and in response to copper handling defects in other tissues. Intracellular copper is trafficked by a series of dedicated chaperones that form an effective delivery pathway to target enzymes for metalation or labile pools for storage (Cobine et al., 2021). Many of the central factors for copper transport have been elucidated and disruptions to these steps can exert devastating effects on human health. In our mouse models of mitochondrial dysfunction, we observe a signature of disrupted copper homeostasis in the liver and a responsive cascade of subsequent compensatory events. This response initiates with the progressive loss of the CTR1 copper transporter in the liver via proteasomal degradation, effectively reducing copper uptake (Hlynialuk et al., 2015). However, a shift to enhance copper efflux from the tissue likely follows, given that the levels of copper and the copper-dependent ferroxidase ceruloplasmin are both elevated in *hep* plasma (Hlynialuk et al., 2015). Consistent with an important role for hepatic copper mobilization in the observed immune phenotypes, the liver of *Ctr1*^*hrt*^ mice upregulates ATP7A expression and pumps copper into the circulation as a consequence of loss of CTR1 function in the heart, a response that renders it copper-deficient relative to the liver of wild-type littermates (Kim et al., 2010).

To reconcile our findings, we propose the following model. The induction of AFP synthesis in response to mitochondrial dysfunction coincides with the cellular response to increased copper efflux into the secretory system. Although the exact role of copper in potentiating the immunosuppressive properties of AFP is currently unclear, we speculate that AFP is copper-loaded prior to its secretion. Our model accounts for the apparent inert activity of high circulating levels of AFP in the early post-natal period and makes clear predictions that disruptions to copper efflux into the secretory system would ablate the leukocyte toxicity. Copper delivery to the secretory system is mediated via ATOX1 to the ATPase copper pumps, ATP7A and ATP7B (Hamza et al., 2003). Pathogenic variants in *ATP7A* and *ATP7B* manifest as Menkes and Wilson disease, respectively (Bull et al., 1993, Vulpe et al., 1993). In line with our model, neither disorder is characterized by immune suppression or neutropenia, even though severe liver pathology is a hallmark of Wilson disease (Schilsky, 2014). Mitochondrial signaling has previously been shown to promote cellular copper efflux (Leary et al., 2007, Leary et al., 2013), and the most severe leukopenias in our *hep* models coincide with higher levels of copper in the circulation. While these findings support the notion that sufficient ATP7A and ATP7B remains localized within the secretory system to account for the observed phenotypes in liver experiencing mitochondrial dysfunction, future investigations will be required to untangle the trafficking dynamics of the efflux transporters in hepatocytes under these conditions.

A key question remains, which is how is the transcriptional suppression of AFP expression released? Although we observe ISR activation upon mitochondrial dysfunction in the liver, it may be that it is not the signaling cascade that regulates AFP induction. ISR activation is emerging as a robust stress response to mitochondrial OXPHOS dysfunction in patients and in animal models of these diseases (Davis et al., 2016, Koene et al., 2015, Lehtonen et al., 2016, Mick et al., 2020, Yatsuga et al., 2015). The signaling cascade leads to the induction and secretion of two growth factors, GDF15 and FGF21, that remodel cellular metabolism in response to nutritional and metabolic disturbances. Thus, while induction of this stress response is not surprising in our mouse models, ISR activation in other mitochondrial disorders caused by OXPHOS dysfunction is not always accompanied by increased AFP expression or a leukopenia(Forsstrom et al., 2019, Kuhl et al., 2017). We propose instead that the central role of mitochondrial signaling in regulating copper homeostasis in the liver likely underpins the mechanism for AFP immunosuppression. The basis of this pathway opens up an exciting new avenue for the role of mitochondria in intracellular signaling.

In conclusion, we identify a novel tissue crosstalk mechanism whereby the homeostatic role of mitochondria in coordinating hepatic copper homeostasis intersects with the function of the peripheral immune system. Diet could also be a key factor in modulating the response, particularly with foods rich in copper. Future studies will explore the role of this leukopenia mechanism in the underlying susceptibility of patients with mitochondrial disease or ataxia telangiectasia to infections.

## Supporting information

Mass spectrometry data

## Acknowledgements

The authors are indebted to Drs. Dennis J. Thiele (Sisu Pharma), Eric A. Shoubridge (McGill University), Dennis Winge (Utah University) and Kerry Lavender and Scott Wiedenmaier (University of Saskatchewan) for helpful discussions. This work was supported by grants-in-aid of research from the Canadian Institutes for Health Research (MOP16652, S.C.L.), National Institutes of Health (DK110195, B.-E.K.; GM120211, P.A.C and S.C.L; GM111672, V.M.G) and Barth Syndrome Foundation (V.M.G). PAC is supported by the Alabama Agricultural Experiment Station,

## Author contributions

SCL and KAJ conceived the original study, while KAJ, PAC, B-EK, H-YMC, VMG and SCL contributed to its further refinement. Data were collected by KAJ, ZNB, AB, MAH, OY, PAC, SG, KB, SY and CL, and analyzed by KAJ, SG, KB, CL and SY. JD, PN, KM, CS and GNI provided guidance, reagents or samples invaluable to experimental advancement. The manuscript was written by SCL, BJB and KAJ and edited by PAC, BJB, B-EK, H-YMC and VMG.

