## Supplementary methods for "Mitochondrial dysfunction triggers secretion of the immunosuppressive factor α-fetoprotein"

**STAR METHODS**

**KEY RESOURCES TABLE**

| Reagent or Resource | Source | Identifier |
|---|---|---|
| **Antibodies** | | |
| SCO1 | Rabbit polyclonal | In-house |
| GAPDH | Cell Signaling | 2118 |
| ATP7A | In-house | (Hlynialuk et al., 2015) |
| CTR1 | GenScript | In-house |
| eIF2α | Cell Signaling | 9722 |
| phospho-eIF2α | Cell Signaling | 9721 |
| Actin hFAB rhodamine | Cell Signaling | 12004163 |
| SD70 | Mitoscience | MS204 |
| Core 1 | Mitoscience | MS303 |
| ATP5A | Abcam | Ab14748 |
| NDUFA9 | Mitoscience | MS111 |
| COX1 | Mitoscience | MS404 |
| COX4 | Mitoscience | MS407 |
| TOM40 | Proteintech | 18409-1-AP |
| AFP | Abcam | Ab46799 |
| Tubulin | Santa Cruz | sc-5286 |
| CD16/CD32 (Fc Block) | BD Biosciences | 553141 |
| CD3 | BD Biosciences | 565643 |

| | | |
|---|---|---|
| CD4 | BD Biosciences | 557308 |
| CD8 | BD Biosciences | 553030 |
| CD25 | BD Biosciences | 552880 |
| CD44 | BD Biosciences | 563736 |
| CD69 | BD Biosciences | 561240 |
| FITC Annexin V | BD Biosciences | 556419 |
| Annexin V Binding Buffer, 10X | BD Biosciences | 556454 |
| **Bacterial and Virus Strains** | | |
| XL1-Blue Competent Cells | Agilent | 200249 |
| HDΔ28E4LacZ | In-house | (Brunetti-Pierri and Ng, 2011, Brunetti-Pierri et al., 2013) |
| HDΔ24.7E4-pepck-mSCO1 | In-house | (Brunetti-Pierri and Ng, 2011, Brunetti-Pierri et al., 2013) |
| **Chemicals, Peptides, and Recombinant Proteins** | | |
| Ammonium persulfate | BioShop | AMP001.100 |
| Acrylamide | Sigma | A8887-500G |
| Bathocuproinedisulfonic acid disodium salt | Sigma | B1125-1G |

| | | |
|---|---|---|
| Calcium chloride dihydrate | EM Science | B10070-34 |
| Cupric chloride dihydrate | BioShop | CUC002.100 |
| Cupric sulfate pentahydrate | Sigma | C-8027 |
| Dimethyl sulfoxide | BioShop | DMS666.50 |
| Ethylenediamine | BioShop | EDT001.500 |
| Ethylenediamine tetraacetic acid dipotassium salt dihydrate | Fisher | BP19-500 |
| Glycine | BioShop | GLN002.5 |
| L-Histidine | Sigma | H9288 |
| Maraviroc | Sigma | PZ0002 |
| PMSF | BioShop | PMS123.5 |
| Potassium chloride | EM Science | PX1405-1 |
| SDS | BioShop | SDS001.1 |
| Sodium chloride | BioShop | SOD002.5 |
| Potassium phosphate, monobasic | BioShop | PPM302.500 |
| Sodium phosphate, dibasic | BioShop | SPD579.1 |
| Triton-X 100 | BioShop | TRX777.5 |
| Tween-20 | Fisher | BP337-500 |
| Trypan blue solution (0.4%) | Sigma | T8154 |
| Tris | BioShop | TRS001.5 |
| Skim Milk (Blotto) | US Biological | S1013-90A |
| TEMED | BioShop | TEM001.5 |

| | | |
|---|---|---|
| Complete Protease Inhibitor Cocktail (PIC) | Roche | 11 836 145 001 |
| Hydrogen peroxide solution | Sigma | 216763 |
| Albumin (Bovine) | BioShop | ALB001.500 |
| Dithiotreitol (DTT) | Fisher | BP172-5 |
| Oxaloacetic acid (OAA) | Sigma | O4126-1G |
| Proteinase K from *Tritirschium album* | Sigma | P2308-100MG |
| Cytochrome *c* | Sigma | C-7752 |
| Agarose | BioShop | AGA002.1 |
| SYBR Safe DNA gel stain | Invitrogen | S33102 |
| Generuler (100bp) | Thermo | SM0241 |
| p-Coumaric acid | Sigma | C9008-5G |
| Luminol | Sigma | A8511-5G |
| Ponceau S | Fisher | 40058003755 |
| Recombinant Mouse $\alpha$-fetoprotein (AFP) | Mybiosource | MBS717717 |
| Trichloroacetic acid (TCA) | Sigma | T0699 |
| RPMI 1640 Medium | GE | SH30096.01 |
| 10X PBS | Fisher | BP3994 |
| FBS | Gibco | 12483020 |
| Recombinant human interleukin-2 (IL-2) | Stem Cell | 78036.3 |
| Ficoll Paque Plus | Sigma | 17-1440-02 |
| Hank's Balanced Salt Solution (HBSS) (10X) | Gibco | 14065056 |

| | | |
|---|---|---|
| Isoflurane | Fresenius Kabi | CP0406V2 |
| Harris Hematoxylin | Leica | 3801562 |
| Eosin Y | Leica | 3801602 |
| **Critical Commercial Assays** | | |
| Phire Animal Tissue Direct PCR kit | Thermo | #F-140WH |
| Mouse AFP Quantikine ELISA Kit | R&D Systems | MAFP00 |
| ENLITEN® ATP Assay System Bioluminescence Detection Kit for ATP Measurement | Promega | FF2000 |
| Qproteome Total Glycoprotein Kit | Qiagen | 37541 |
| **Experimental Model: Organisms/Strains** | | |
| *Alb-cre* | Jax | #003574 |
| *Sco1* | In-house | (Hlynialuk et al., 2015) |
| *Coa5* | KOMP - ES cells; Toronto Centre for Phenogenomics - mice | |
| *Cox10* | Jax | #024697 |
| *Ctr1* | In-house | (Kim et al., 2010) |
| **Oligonucleotides** | | |

| | | |
|---|---|---|
| Sco1-F<br><br>ATGGAATCCCTTCCTTGCTTC | In-house | (Hlynialuk et al., 2015) |
| Sco1-R1<br><br>TCAACCTCAACATTTCACGACGGTATT | In-house | (Hlynialuk et al., 2015) |
| Sco1-R2<br><br>ACCTAAAAGTGGGGCTTCCTGAAAACTAA | In-house | (Hlynialuk et al., 2015) |
| Cox10-F<br><br>GAGAGGAGTCAAGGGGACCT | This study | N/A |
| Cox10-R1<br><br>GGCCTGCAGCTCAAAGTGTA | This study | N/A |
| Cox10-R2<br><br>CAAAGAGGGCTCACTTCTTGC | This study | N/A |
| Coa5-F<br><br>GAGCTCTCATGCACAGCAAG | This study | N/A |
| Coa5-R1<br><br>TTCAAGTCGTGGAATGGTAGC | This study | N/A |
| Coa5-R2<br><br>GCTGCTAGGACCAAATCCTG | This study | N/A |
| Ctr1-P1<br><br>AATGTCCTGGTGCGTCTGAAA | In-house | (Kim et al., 2010) |
| Ctr1-P2<br><br>GCAGTAGATAAAAGCCAAGGC-30 | In-house | (Kim et al., 2010) |

| | | |
|---|---|---|
| Ctr1-P3<br><br>AAAAACCACTATTCAGAGACTG | In-house | (Kim et al.,<br><br>2010) |
| Afp-F<br><br>AGTTGCAAAGCACATGAAGA | IDT | |
| Afp-R<br><br>AAGCACTCCTCCTTGTTGTC | IDT | |
| Gapdh-F<br><br>CATGGCCTTCCGTGTTCCTA | IDT | |
| Gapdh-R<br><br>CCTGCTTCACCACCTTCTTGA | IDT | |
| **Software & Algorithms** | | |
| GraphPad Prism 8.0/9.0 | | |
| CytExpert | | |
| ImageJ | | |

### LEAD CONTACT AND MATERIALS AVAILABILITY

Information and requests for resources and reagents should be directed to and will be fulfilled by the Lead Contact, Dr. Scot C. Leary. This study did not generate any new, unique reagents.

### EXPERIMENTAL MODEL AND SUBJECT DETAILS

*Animal models and husbandry*

Homozygous floxed *Sco1*, *Cox10* and *Coa5* mice were used to generate hepatocyte-specific (*hep*) knockout models as previously described (Hlynialuk et al., 2015). Briefly, floxed animals were crossed with mice in which *Cre* recombinase expression is driven by the albumin promoter (*Alb-Cre*$^{tg/tg}$). *Cre* positive, F1 progeny were then backcrossed to the appropriate homozygous floxed model to generate F2 litters, with roughly 25% of the resultant progeny exhibiting hepatocyte-specific loss of expression of the gene of interest.

Heart-specific *Ctr1* (*Ctr1*$^{hrt}$) knockout mice were generated according to (Kim et al., 2010). PCR genotyping of *Sco1* (Hlynialuk et al., 2015), *Cox10* (Diaz et al., 2008) and *Ctr1* (Kim et al., 2010) mice was as previously described, and age-matched *flox/+* or *flox/flox* siblings served as *Controls* in this study. *Coa5* mice were genotyped as detailed below. Mice were housed under a 12hr light: 12hr dark photoperiod in a temperature and humidity-controlled facility and provided with food and water *ad libitum*. All experiments on F2 *hep* animal models were approved by the Animal Ethics Review Board at the University of Saskatchewan (AUP# 20100091, S.C.L.) while those involving the *Ctr1* mouse model were conducted in accordance with National Institutes of Health Guide and approved by the Institutional Animal Care and Use Committee at the University of Maryland, College Park (AUP# R-APR-18-14, B.-E.K).

*Generation of a conditional Coa5 mouse model*

ES cells with floxed *Coa5* alleles were purchased from the Knockout Mouse Project (KOMP) Repository. Male chimera transmitter *Coa5* mice lacking the neomycin-resistance cassette and *lacZ* reporter gene were then generated fee-for-service at the Toronto Centre for Phenogenomics and shipped to the University of Saskatchewan. These males were crossed with C57BL/6N females, and the resultant *Coa5*$^{loxP/wt}$

progeny were intercrossed to yield homozygous *Coa5* mice with a residual FRT site and loxP sites that flank the second exon of the gene to allow for its deletion.

### METHOD DETAILS

#### *Specimen collection*

Mice were weighed prior to being anesthetized with 2% isofluorane. Blood was then collected from the left ventricle of the heart using a 27-gauge needle, placed into a precoated $K_2$EDTA tube, and inverted 10 times. After a 20 minute room temperature incubation, 100μl of blood was retained on ice for further analyses while the remainder was used to isolate plasma by two sequential room temperature spins for 5 minutes at 1,000 *x g*.

Following blood collection, the thymus, heart, spleen, liver, kidney and brain were harvested, and flash frozen on dry ice in pre-weighed Eppendorf tubes. Tubes were then weighed post-collection, allowing for tissue wet weights to be calculated. Tissues were then powdered in a stainless-steel mortar and pestle on dry ice and stored at -80$^{o}$C for subsequent biochemistry, molecular biology, elemental and genetic analyses.

#### *Complete blood cell count (CBC) and peripheral blood smear*

Automated CBC analysis was performed on the fraction of retained blood using the COULTER® Ac·T diff™ Analyzer (Beckman). All samples were run in duplicate. Alternatively, a spreader slide and 10μl of blood were used to create a peripheral blood smear. Air-dried slides were then stained with Wright's Geimsa using an immersion protocol. Briefly, slides were stained for 1 minute, rinsed for 5 min in phosphate buffer (pH 6.8) (made with potassium phosphate, monobasic 50.1% (w/w) and sodium phosphate, dibasic 49.9% (w/w)), washed briefly in running deionized water, dried and cover slipped. The white blood cell (WBC) count was determined by taking the average number of WBCs in 10 fields at 40X high power and multiplying by $2.0x10^9$/L (Jones, 2009).

*Tissue perfusion, fixation and histology*

Mice were anesthetized with 2-3% isoflurane and oxygen, and perfused through the heart with 10 ml HBSS (pH 7.2) containing 2.5% FBS at a flow rate of ~5ml/min. Once the organs were cleared of blood, mice were perfused with another 10 ml of 10% formalin. The thymus, heart, spleen and liver were harvested, stored overnight in 10% formalin, then dehydrated and embedded in paraffin. Seven micrometer cross-sections were prepared, and slides were stained with hematoxylin and eosin (H&E) according to the manufacturer's standard procedure. Sections were viewed and imaged using a scanning transmission light microscope (Leica).

*Adenoviral-related experiments*

Helper-dependent adenovirus (HdAD) encoding the *lacZ* transgene or vehicle (sterile Ringer's solution) were administered via intracardiac (IC) or interperitoneal (IP) injection. We found that IC administration resulted in higher β-galactosidase expression in the liver with less intense staining in other, peripheral tissues when compared to IP injection (Fig. S1B). We therefore used the IC route to administer $5.783x10^{12}$ *Sco1* HdAD particles/kg or an equivalent volume of vehicle to mice at 21-24 days of age. Mice were anesthetized with 2% isofluorane, the chest and abdomen were disinfected with 70% ethanol, and adenovirus or vehicle was administered with a 27-gauge needle by direct puncture of the left ventricle through the diaphragm following the drawback of fresh arterial blood. Blood and tissues were then collected when mice reached 47 days of age.

*Immunoblot analyses*

Powdered whole tissues were resuspended and homogenized on ice in extraction buffer (Potting et al., 2013) supplemented with a complete protease inhibitor cocktail (Roche) and 0.5mM PMSF. Following a 30 min incubation step on ice, lysates were clarified by centrifugation at 16,600 *x g* for 10 minutes at 4$^{o}$C and equal amounts of protein (10-25μg/lane) were electrophoresed on 4-20% pre-cast Tris-HCl gels (BioRad). All gels were then transferred onto nitrocellulose membrane under semi-dry conditions and blotted with the appropriate antibodies. Following incubation of membranes with secondary antibody, immunoreactive proteins were detected by luminol enhanced chemiluminescence. For cultured cells, 2 X 10^5 cells were seeded on a 6 well plate and treated with either DMSO or OXPHOS inhibitors including Rotenone (1 μM), Malonate (1 mM), Antimycin (1 μM), KCN (1 mM), Oligomycin (1 μM) for 24 hours. Cells were washed with PBS and harvested using a cell scraper. Cells were then lysed in RIPA buffer supplemented with protease inhibitor cocktail (Roche). After 30 min incubation in lysis buffer, cell lysates were collected by centrifugation at 14, 000 *x g* for 10 mins at 4˚C and equal amounts of protein (20 μg) were electrophoresed on 10% pre-cast Bis-Tris gels (Life Technologies). Following semi-dry transfer of gels onto PVDF membranes, the membranes were blotted with indicated antibodies (anti-AFP obtained from Leary Lab and anti-β-actin from Sigma A2228) and protein bands detected as above.

*Elemental analyses*

Samples were digested in 40% nitric acid by boiling for 1 hour in capped, acid washed tubes, diluted in ultra pure, metal free water and analyzed by ICP-OES (Perkin Elmer, Optima 7300DV) versus acid washed blanks. Concentrations were determined from a standard curve constructed with serial dilutions of two commercially available mixed metal standards (Optima). Blanks of nitric acid with and without “metal-spikes” were analyzed to ensure reproducibility.

*qPCR analyses*

Total RNA was extracted with Trizol Reagent and quantified using a Nanodrop. cDNA was synthesized using Superscript IV Reverse Transcriptase and Oligo(dT)20 primers from ThermoFisher. qPCR was performed using SsoFast EvaGreen Supermix (Bio-Rad) on a CFX384 Touch Real-Time PCR Detection System (Bio-Rad). Data were analyzed according to the delta delta Ct method and *Afp* levels were normalized against *Gadph* abundance.

*ATP quantitation*

ATP levels in $Sco1^{hep}$, $Cox10^{hep}$, $Coa5^{hep}$ and age-matched littermate *Control* livers were measured using a luminescent detection kit, as per the manufacturer's instructions. Briefly, 10mg of powdered liver tissue was homogenized in 2.5% trichloroacetic acid (TCA), sonicated on ice and neutralized with Tris-acetate buffer. Signal intensities were read using a Luminex multimode plate reader. Arbitrary luminescence units in *hep* livers were represented as the fold change relative to the appropriate *Control* group.

*Tail vein injections*

Plasma for tail vein injections was aseptically pooled, aliquoted and stored at -80$^{o}$C prior to the beginning of an experiment. *Control* mice were injected via the tail vein between 24 and 27 days of age with a 27-gauge needle, after tails had been warmed for 5-10 min under a heat lamp and subsequently cleaned with 70% ethanol to prevent infection. Animals were injected with the maximum recommended volume of 10ml/kg or 100-200μl of plasma isolated from *Control* or *Sco1*$^{hep}$ mice, or with 1μg of rAFP or albumin, a closely related family member. Mice were injected every 72 hours thereafter for a 28 day period. Blood and tissues were collected 3 days after the eighth injection.

*PBMC isolation and culture*

Mouse blood was collected as described above. Human blood was obtained from three consenting, healthy volunteers (30-50 years of age) and used in experiments approved by the Research Ethics Board at the University of Saskatchewan. Both mouse and human PBMCs were isolated by Ficoll Plaque Plus density centrifugation, and cultured at a concentration of $10^6$ cells/ml in RPMI 1640 media supplemented with 20% FBS and

100 IU IL-2. All related experiments were carried out one to three days after PBMCs were initially isolated.

*Plasma fractionation and PBMC treatment*

Plasma pools were generated for *Sco1*$^{hep}$, *Cox10*$^{hep}$ and age-matched littermate *Control* animals. Plasma from mice fed a high fat (HF) diet (Savard et al., 2013) was also obtained, pooled and used as an additional control. PBMCs were cultured as described above in media containing 20% FBS or 17.5% FBS and 2.5% *Control*, *hep* or *HF* plasma.

To determine if the bioactive factor in *hep* plasma was a protein and the nature of that protein, plasma from *Control* and *Sco1*$^{hep}$ mice was first boiled for 5 min at 98°C or treated with trypsin for 1 hr. PBMCs were then cultured as described above in media containing 17.5% FBS plus 2.5% untreated, boiled or trypsinized plasma from *Control* or *Sco1*$^{hep}$ mice. Control and *hep* plasma was then separated by size using centrifugal 50KDa Filters (Amicon) and by glycan content using the Qproteome Total Glycoprotein Kit, with both steps being done according to the manufacturer's instructions. PBMCs were cultured as described above in media supplemented with 17.5% FBS containing 2.5% of each plasma fraction.

For AFP depletion experiments, a volume of plasma equivalent to 2.5% v/v from *Control* or *Sco1*$^{hep}$ mice was incubated with 1μg mouse anti-AFP and rotated at 4°C overnight. 250μl of protein A dynabeads was then added to the plasma and incubated at room temperature for 2 hours. Mouse anti-SLC25A3 was used as an IgG isotype control. Antibody-bead complexes were collected on a magnet, and the resultant plasma was used to treat PBMCs as described above. *Control* and *Sco1*$^{hep}$ plasma that had been incubated with an equivalent volume of PBS served as internal controls.

*Mass Spectroscopy (MS) and differential analysis*

Quantitative MS analyses were conducted fee-for-service by the Proteomics platform at the McGill University Health Centre. Briefly, 2μg of the >50kDa glycosylated plasma fraction from *Control*, *Sco1*$^{hep}$, *Cox10*$^{hep}$ and *HF* mice was trypsinized and analyzed using a Thermo Scientific Ultimate 3000 HPLC and Orbitrap Fusion MS: Quadrupole-Orbitrap-Linear ion trap hybrid. Raw data was then mined with Pinnacle (http://www.optystech.com/index.html#) and the spectral counts for each peptide were quantified. Only proteins for which two unique peptides were detected were included in the final analyses, and quantitative data for each experimental group represent the average of two independent MS runs.

Liquid chromatography MS (LCMS) differential analysis was performed using R (R Core Team, 2017). The package DESEQ2 (Love et al., 2014) was used to test for differential expression, the package data.table (Dowle and Srinivasan, 2019) was used for data manipulation and ggplot2 was used to generate plots (Wickham, 2016). Raw data was imported (20190508_LCMS_raw_data_sheet1) and counts were converted to integers. Counts with NA were converted to zero. One count was added to all samples. The count matrix was prepared (20190508_input_sheet2) and DESeq2 was performed on *hep* vs *Control*. A false discovery rate (FDR) of 0.1 was applied to report significant (upregulated/downregulated) versus non-significant (ns) hits (20190508_results_sheet3).

*Flow Cytometry*

PBMCs isolated from *Control* and *Sco1$^{hep}$* blood was stained with an anti-CD45, CD3, CD4, CD25, CD44 and CD69 antibody cocktail or with an anti-CD3, CD4, 7AAD and Annexin V antibody mixture. Briefly, PBMCs were resuspended in PBS with 2.5% FBS at $1\times10^6$ cells/mL. Cells were blocked with anti-mouse CD16/CD32 Fc block for 15 min and then incubated with the primary antibody cocktail for 30 min. For cells stained with Annexin V, PBMCs were resuspended and blocked as described above, and then incubated with anti-CD3 and anti-CD4 for 20 min. 7AAD antibody was subsequently added to the staining mixture and further incubated for 10 min. Cells were then centrifuged for 5 min at 1000 *x g*, resuspended in Annexin V binding buffer and stained according to the manufacturer's instructions. Following either staining procedure, cells were fixed in a solution of 1% formalin with red cell lysing solution for 15 min. All events were analyzed using a CellFlex™ Flow Cytometer (Beckman Coutler). Dead cells were excluded based on forward and side light scattering. All antibody staining procedures took place at room temperature in the dark.

### QUANTIFICATION AND STATISTICAL ANALYSIS

All statistical analysis was performed using GraphPad Prism 8.0 or 9.0. Data were reported as mean ± SEM. For selection of appropriate statistical tests, data were assessed by histogram or Tukey plot to detect a normal Gaussian distribution. After verifying a Gaussian distribution, statistical differences between two groups were assessed using a Student's t-test and between three or more groups with a one-way

ANOVA and a Tukey's or Sidak post-hoc test. For the qPCR analyses in Figure S5, differences in *Afp* mRNA abundance between the *Control* and $Ctr1^{hrt}$ hearts were assessed using a non-parametric Mann-Whitney U test as the data were not normally distributed. For data with two independent variables, a two-way ANOVA and Bonferonni's post-hoc test was used. p-values <0.05 were considered significant. Statistical parameters can be found in the figure legends.

### DATA AND SOFTWARE AVAILABILITY

The datasets supporting the current study have not been deposited in a public repository but are available from the corresponding author on request.
