## Supplementary Figures for "Mitochondrial dysfunction triggers secretion of the immunosuppressive factor α-fetoprotein"

**Supplemental Data**

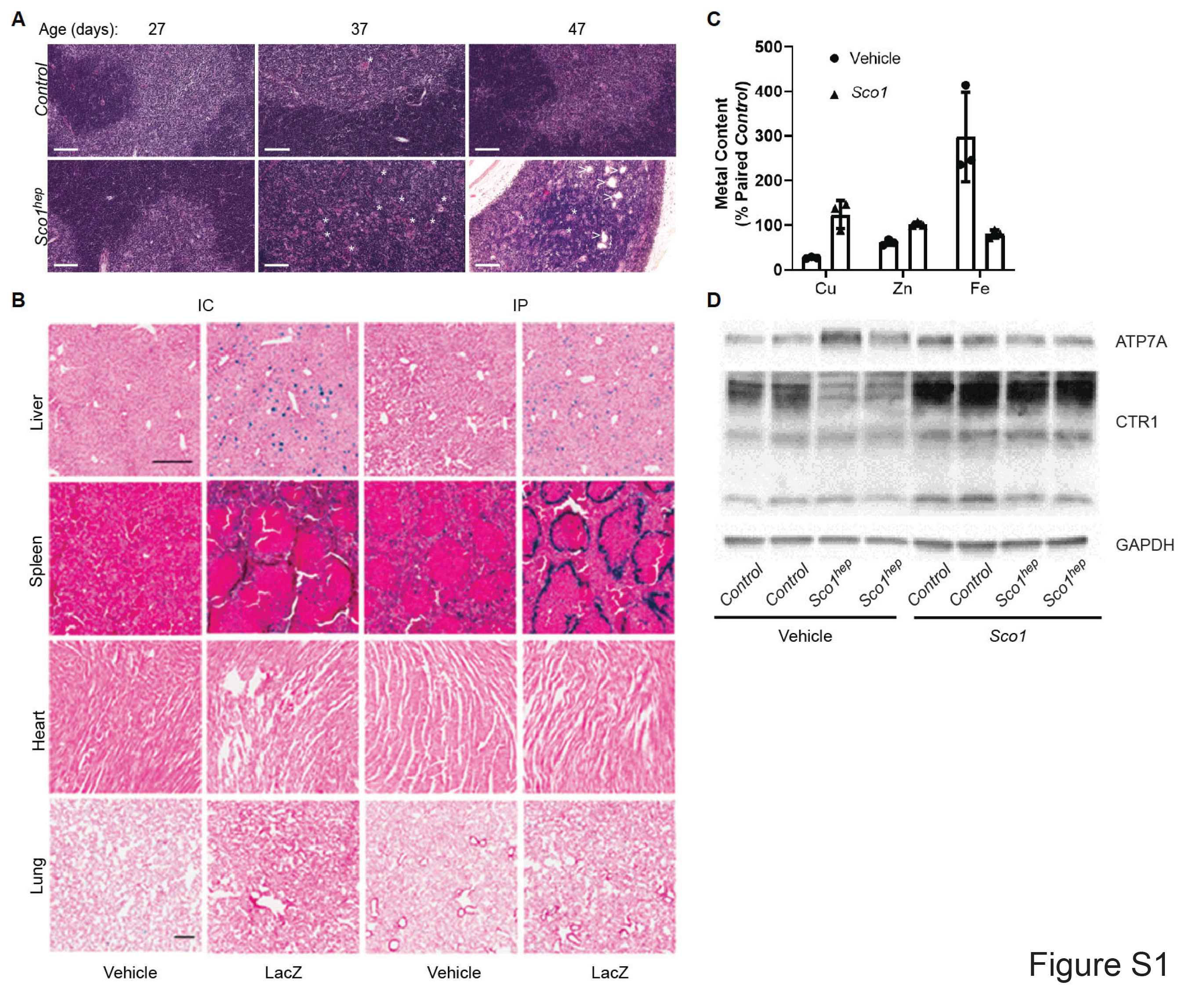

**Figure S1.** *A)* The *Sco1^hep^* thymus exhibits progressive thinning of the cortex from P27 to P47, with disruption of the cortico-medullary boundary, accumulation of tingible body macrophages (denoted with a *) and increased vascularity (denoted with a >). Scale bar, 100μm. *B)* LacZ staining in the liver (4X), spleen (4X), heart (4X) and lung (2X) upon intracardiac (I*C*) or intraperitoneal (IP) administration of vehicle or helper-dependent adenovirus. *C&D)* Restoration of *Sco1* expression in the *Sco1^hep^* liver normalizes *C)* metal ion levels (Cu and Fe, p< 0.01; Zn, p< 0.05) and *D)* CTR1 abundance. *Control* refers to wild-type littermates.

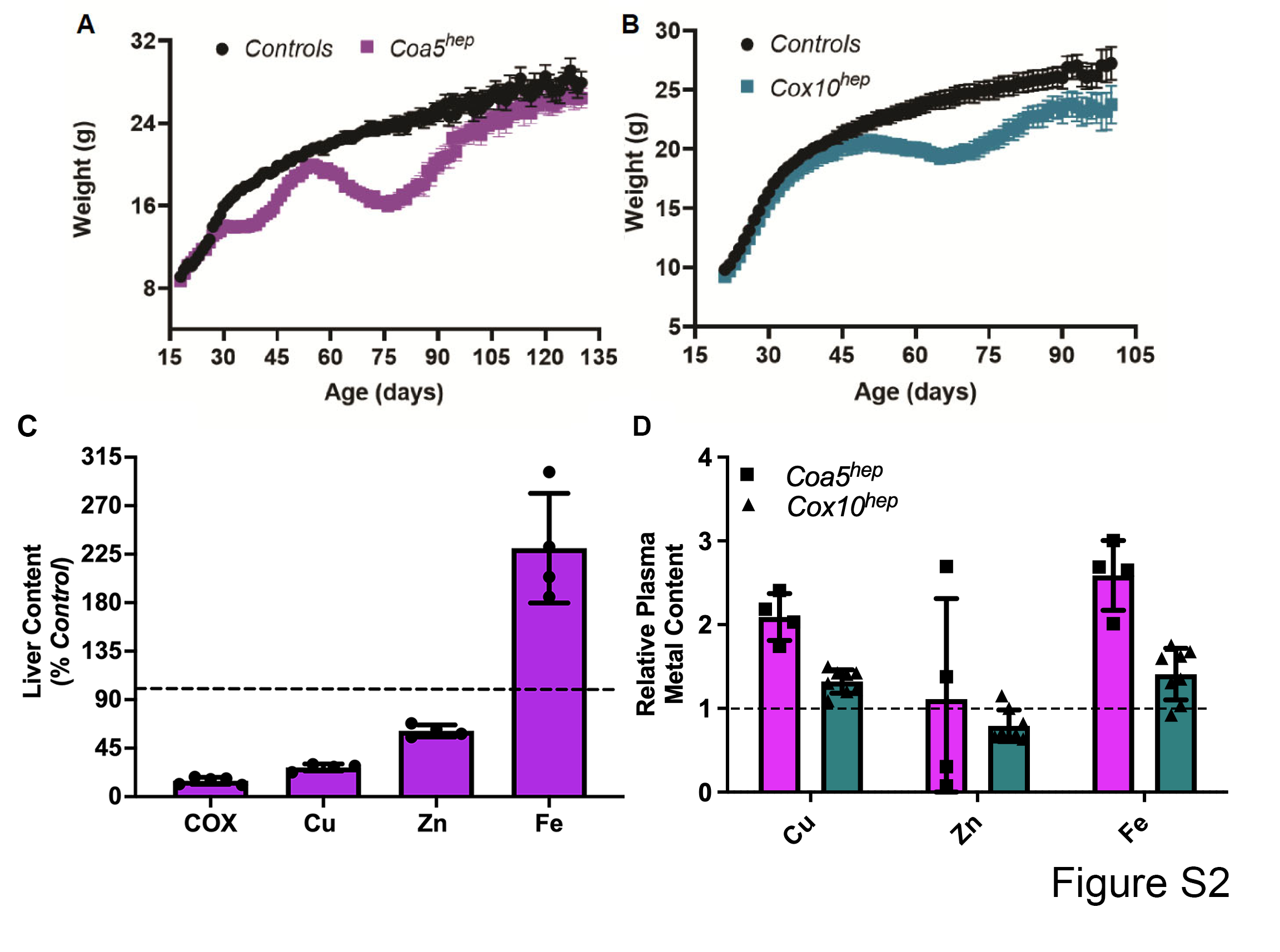

**Figure S2.** Change in body weight (g*)* over time in *A)* *Coa5^hep^* and *B)* *Cox10^hep^* mice. *C)* *Coa5^hep^* livers have a severe COX and copper deficiency relative to livers from *Control* littermates. *D)* Plasma copper, iron and zinc levels in *Coa5^hep^* and *Cox10^hep^* plasma relative to age-matched, littermate *Controls*.

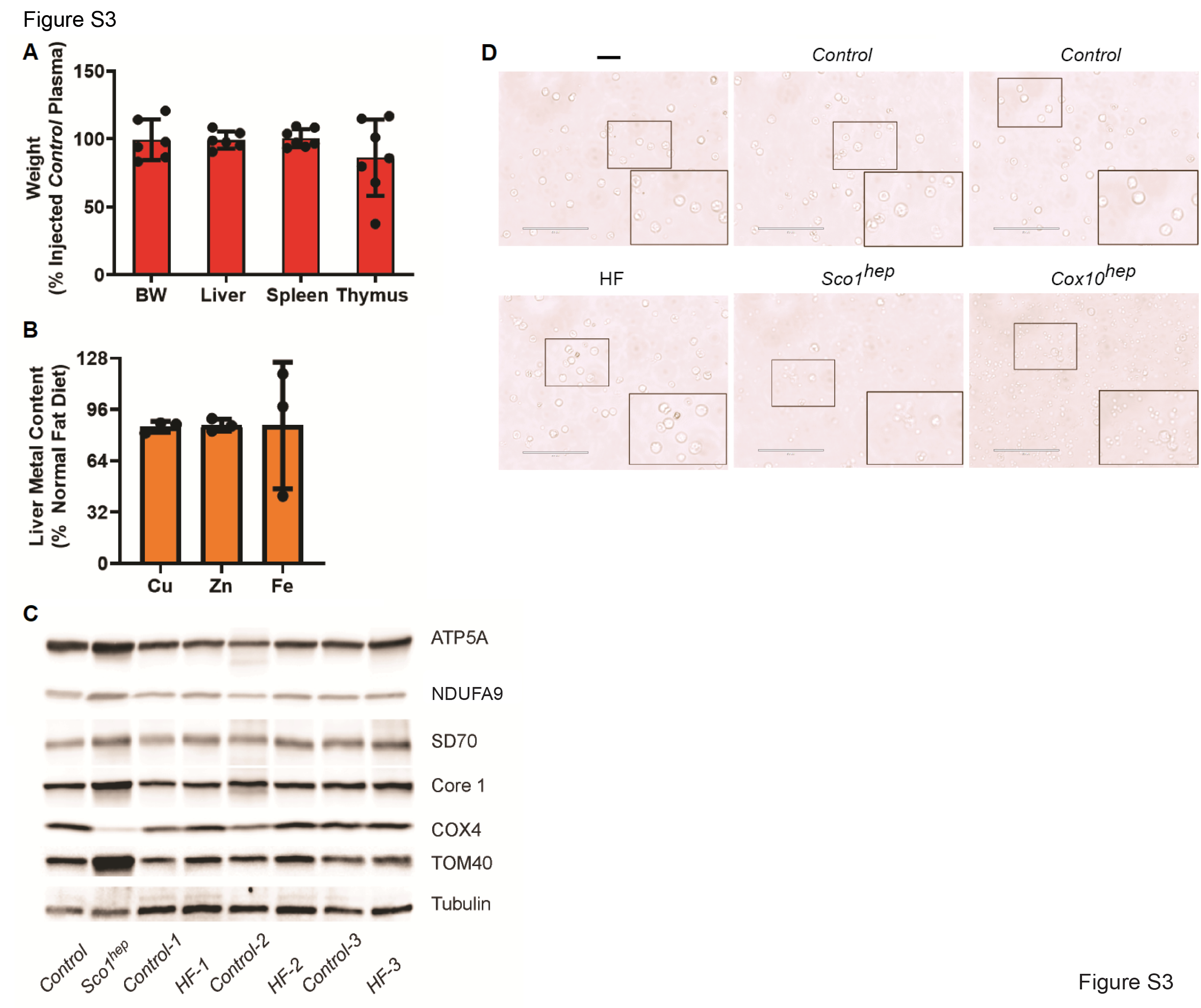

**Figure S3.** *A)* Body and organ weight are unaffected in *Control* mice injected with *Sco1^hep^* plasma relative to those injected with *Control* plasma. *B)* Metal content and *C)* OXPHOS subunit abundance are unaltered in livers from mice fed a high fat (HF) diet compared to those fed normal chow. *Control* and *Sco1^hep^* liver extracts were included for comparative purposes and tubulin served as an internal loading control. *D)* Lower magnification showing a greater number of PBMCs, with black boxes depicting the region of interest shown in Figure 3B.

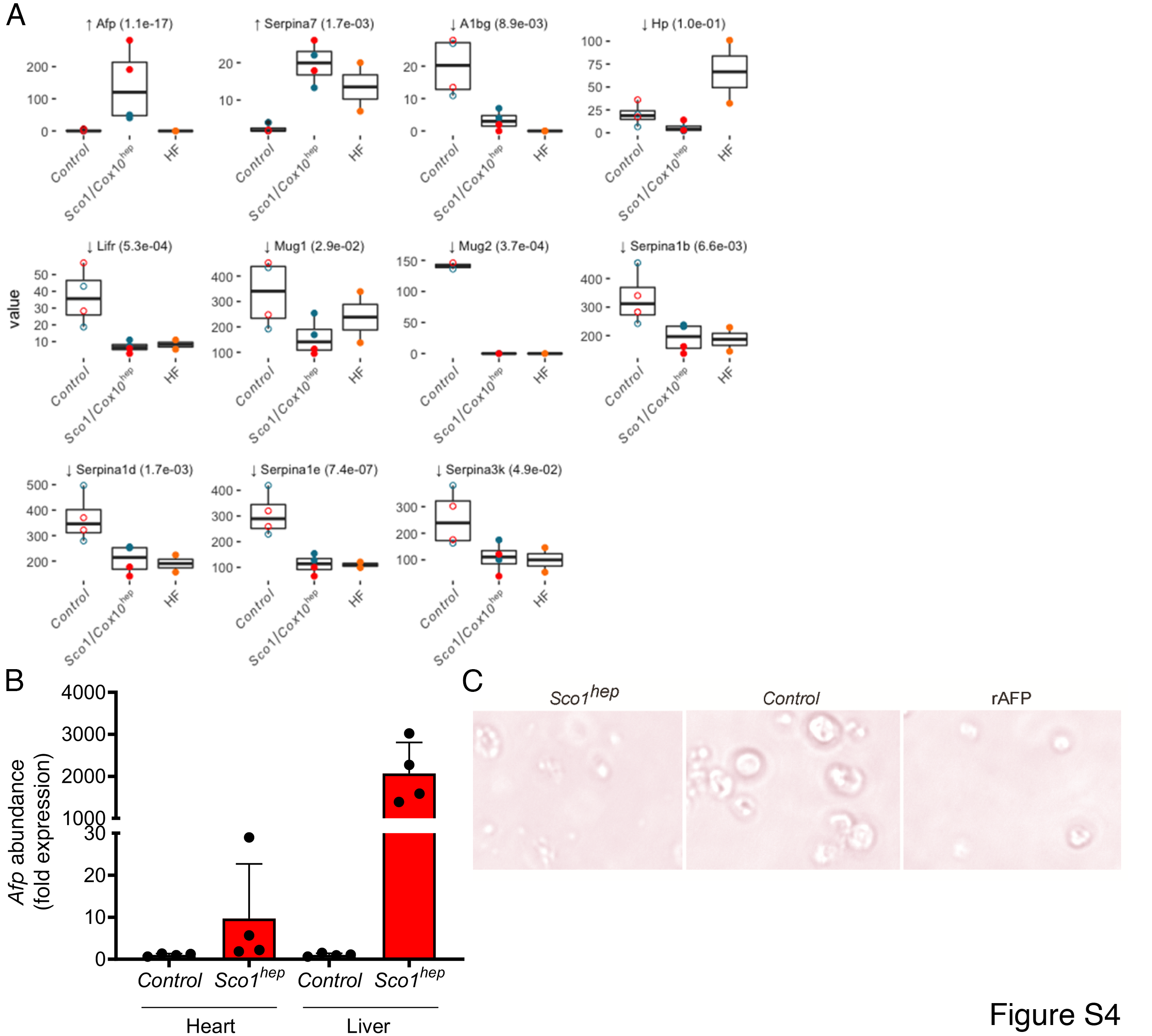

**Figure S4.** *A)* Individual box plots of significantly up- or downregulated plasma proteins in *hep* compared to *Control* mice. Red and blue circles denote data from the *Sco1* and *Cox10* models, respectively (open circles, *Control* animals; closed circles, *hep* animals). HF denotes plasma from mice fed a high fat diet. *B)* *Afp* mRNA levels are significantly higher in the *Sco1^hep^* liver (*p* < 0.02) but not the heart, when compared to *Control* tissues from age-matched littermates. Transcript levels were normalized to *Gapdh* mRNA abundance. *C)* PBMC viability is similarly reduced upon treatment with *Sco1^hep^* plasma or recombinant AFP (rAFP, 1μg).

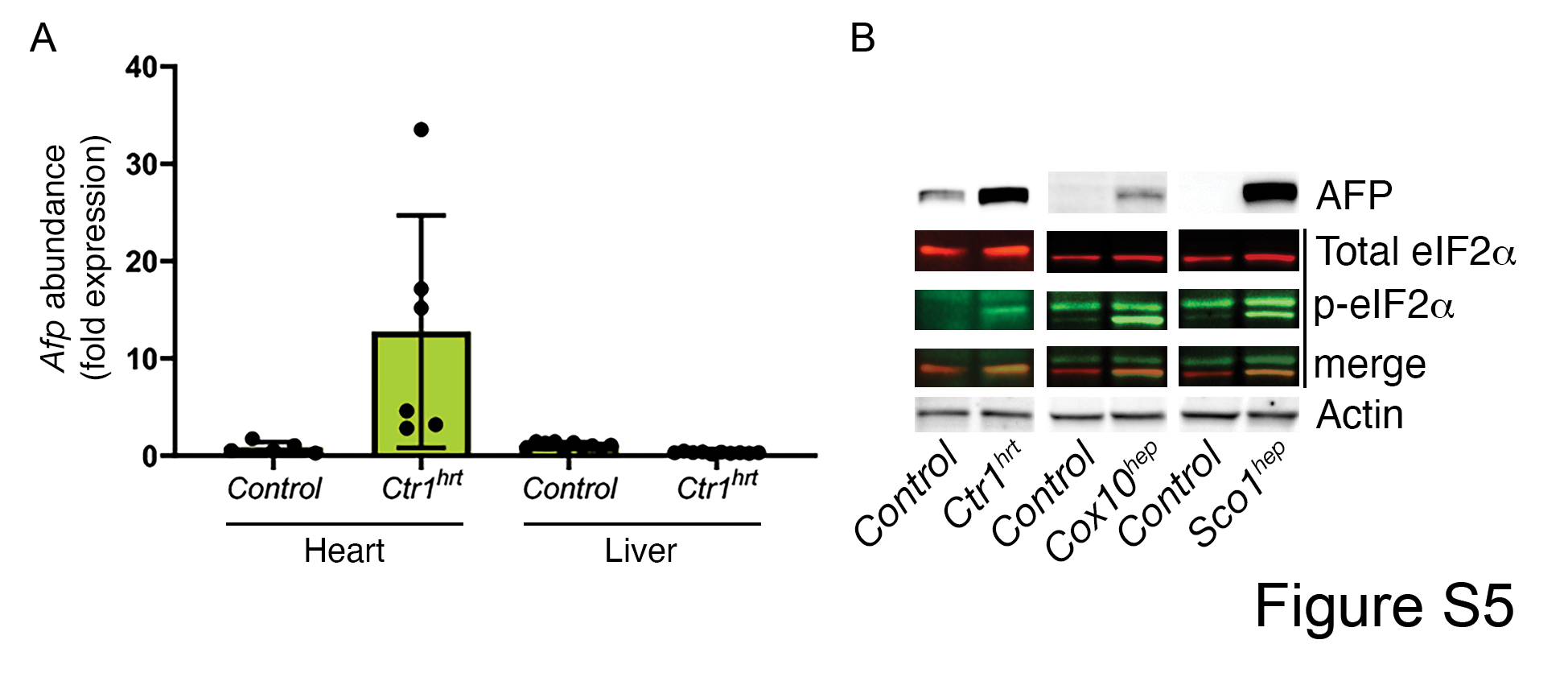

**Figure S5.** *A)* *Afp* mRNA levels are significantly higher in the *Ctr1^hrt^* heart (*p* < 0.005) but not the liver, when compared to *Control* tissues from age-matched littermates. Transcript levels were normalized to *Gapdh* mRNA abundance. *B)* The *Ctr1^hrt^* heart has elevated levels of the ISR marker phospho-eIF2α. Equal amounts of *Control* and *hep* liver extracts from the *Sco1* and Cox10 lines were included in these analyses for comparative purposes.

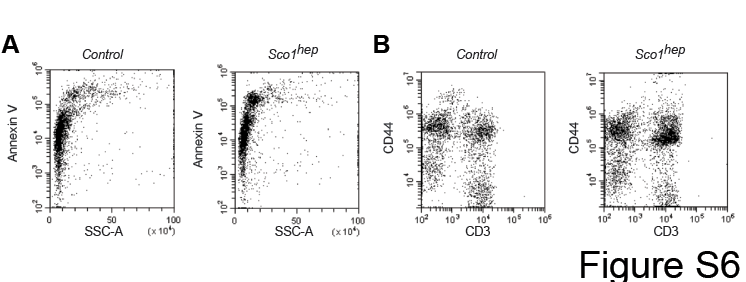

**Figure S6.** Flow plots of peripheral PBMCs show that *Sco1^hep^* mice show have a higher percentage of cells positive for the cell surface expression of *A)* the activation marker CD44 and *B)* the apoptotic marker Annexin 5.
